# Controlled microtrauma opens a regenerative window for appendage-bearing skin repair

**DOI:** 10.64898/2026.06.18.732512

**Authors:** Kang Wang, Zi-Yuan Feng, Zhen-Yu Zhang, Qing-Feng Li, Hui-Qi Xie

**Affiliations:** Department of Orthopedic Surgery and Orthopedic Research Institute, Stem Cell and Tissue Engineering Research Center, State Key Laboratory of Biotherapy, West China Hospital, West China School of Medicine, Sichuan University, Chengdu, Sichuan, China; Department of Plastic and Burn Surgery, West China Hospital, West China School of Medicine, Sichuan University, Chengdu, China; Department of Plastic and Reconstructive Surgery, Shanghai Ninth People’s Hospital, Shanghai Jiao Tong University School of Medicine, Shanghai, China

**Author notes:** Correspondence (Hui-Qi Xie); (Qing-Feng Li); (Zhen-Yu Zhang). These authors contributed equally (Kang Wang, Zi-Yuan Feng).

## Abstract

Adult skin normally resolves injury through rapid closure and fibrotic matrix deposition, often at the cost of permanent appendage loss. We tested whether spatially controlled microtrauma could instead serve as a regenerative entry point when paired with temporally coordinated molecular cues. We engineered a hierarchical extracellular-matrix-based microneedle patch that combines rapid local availability of verteporfin, an inhibitor of YAP-associated mechanotransduction, with sustained retinoic-acid delivery to support follicle-regenerative signalling. The microneedle interface was evaluated in full-thickness rabbit ear wounds, which are prone to hypertrophic scarring, and in Bama miniature-pig wounds, whose skin architecture more closely resembles human skin. Across both models, staged dual-cue treatment accelerated wound closure, reduced collagen-dense scar formation and promoted the appearance of hair-bearing tissue and histologically identifiable follicular structures. These findings support a trauma-guided regeneration framework in which controlled microinjury is used not only for delivery but also to open a transient repair niche that can be molecularly redirected toward appendage-bearing skin restoration.

**Importance:** Microneedles are generally treated as minimally invasive delivery devices. Here, the microinjury itself is incorporated into the therapeutic design. The study provides cross-species proof of concept that a patterned injury interface, combined with staged anti-fibrotic and pro-regenerative signalling, can shift wound repair away from fibrotic closure and toward hair-follicle-containing skin. This concise preprint reports the central concept and the rabbit and porcine evidence supporting it; expanded mechanistic and source datasets will be reported separately.

## Introduction

Adult mammalian skin healing is optimized for rapid barrier restoration rather than complete tissue replacement. This response limits infection and fluid loss but frequently produces fibrotic scars and irreversible loss of hair follicles, sebaceous glands and other appendages [1–7]. The problem is especially relevant after surgical injury, burns, traumatic wounds and pathological-scar excision, where tissue damage is unavoidable but current therapies rarely restore appendage-bearing skin.

Wound-induced hair follicle neogenesis demonstrates that adult skin retains latent regenerative capacity [8–13]. However, this response is strongly dependent on wound geometry, species and local mechanics. Excessive mechanical signalling can reinforce fibroblast activation, inflammation and extracellular-matrix stiffening, thereby favouring scar formation over regeneration [14–20]. Inhibition of mechanotransduction, including YAP-associated fibrotic signalling, can preserve regenerative competence and reduce scarring [21–23], but suppressing fibrosis alone does not necessarily provide the morphogenic signals required to rebuild appendages.

Retinoic acid can support epithelial plasticity, dermal papilla activity and Wnt-associated follicle-regenerative programmes [24–26]. We therefore reasoned that a regenerative intervention should combine an early anti-fibrotic cue with a sustained morphogenic cue. To control both injury geometry and cue timing, we developed a hierarchical microneedle patch based on small-intestinal-submucosa-derived extracellular matrix. The patch creates patterned microtrauma, rapidly exposes the wound to verteporfin and maintains local retinoic-acid availability through a protected backing reservoir (Fig. 1).

**Figure 1.**
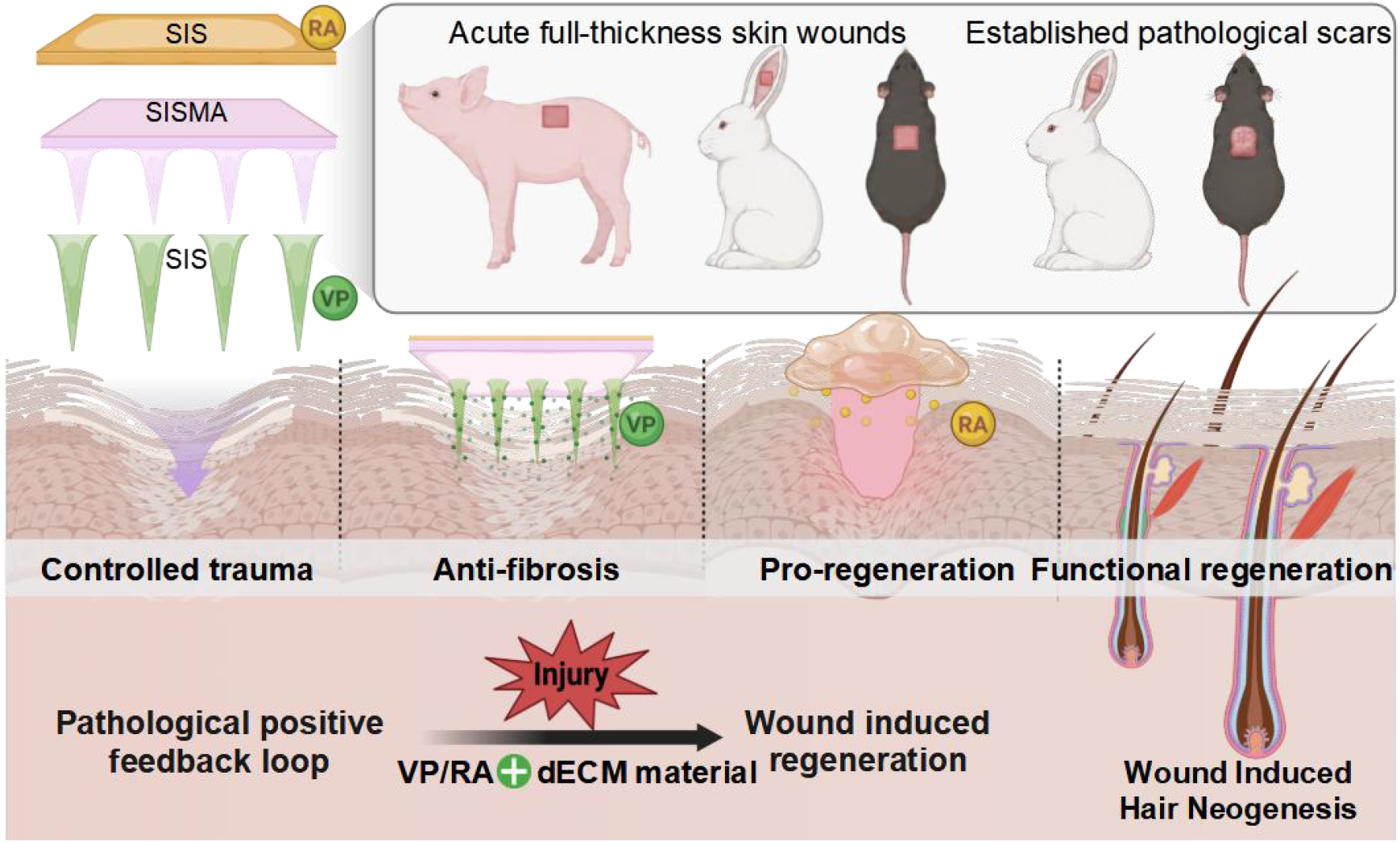
**rauma-guided combinatorial therapy for functional skin regeneration. The hierarchical microneedle patch creates controlled microtrauma and stages local molecular instruction. A verteporfin-containing needle-tip compartment provides an early anti-fibrotic cue, whereas a retinoic-acid-containing backing layer provides sustained pro-regenerative stimulation. The proposed framework is that controlled injury opens a transient repair niche, early restraint of pathological mechanotransduction limits fibrotic reinforcement, and sustained retinoic-acid signalling supports wound-induced follicle regeneration and appendage-bearing repair**.

We focused this concise study on a direct test of the resulting idea in two translationally relevant settings: a rabbit ear model with high scar-forming propensity and a Bama miniature-pig model with human-like dermal architecture. The principal question was whether staged dual-cue treatment could promote hair-follicle-containing repair while reducing fibrotic remodelling across species.

## Results

### A hierarchical microneedle interface couples controlled injury with staged molecular delivery

The patch was constructed as a layered extracellular-matrix interface containing a rapidly dissolving verteporfin-loaded needle-tip component, a photocrosslinked SISMA structural layer and a retinoic-acid-containing backing reservoir. This architecture was selected to match the expected temporal sequence of wound repair: early suppression of mechanically reinforced fibroblast activation followed by sustained support of regenerative signalling. The design also separates the drugs spatially before application and uses microneedle insertion to generate a reproducible array of local microinjuries.

### Staged dual-cue treatment promotes hair-bearing repair in scar-prone rabbit wounds

Full-thickness wounds were created on the ventral rabbit ear, a low-hair-density and high-tension environment prone to hypertrophic scarring. Animals received no microneedle treatment, unloaded microneedles (SM), retinoic-acid-loaded microneedles (R@SM), verteporfin-loaded microneedles (V@SM) or staged dual-loaded microneedles (VR@SM). The VR@SM group showed the fastest macroscopic wound closure among the treatment groups (Fig. 2a–c).

**Figure 2.**
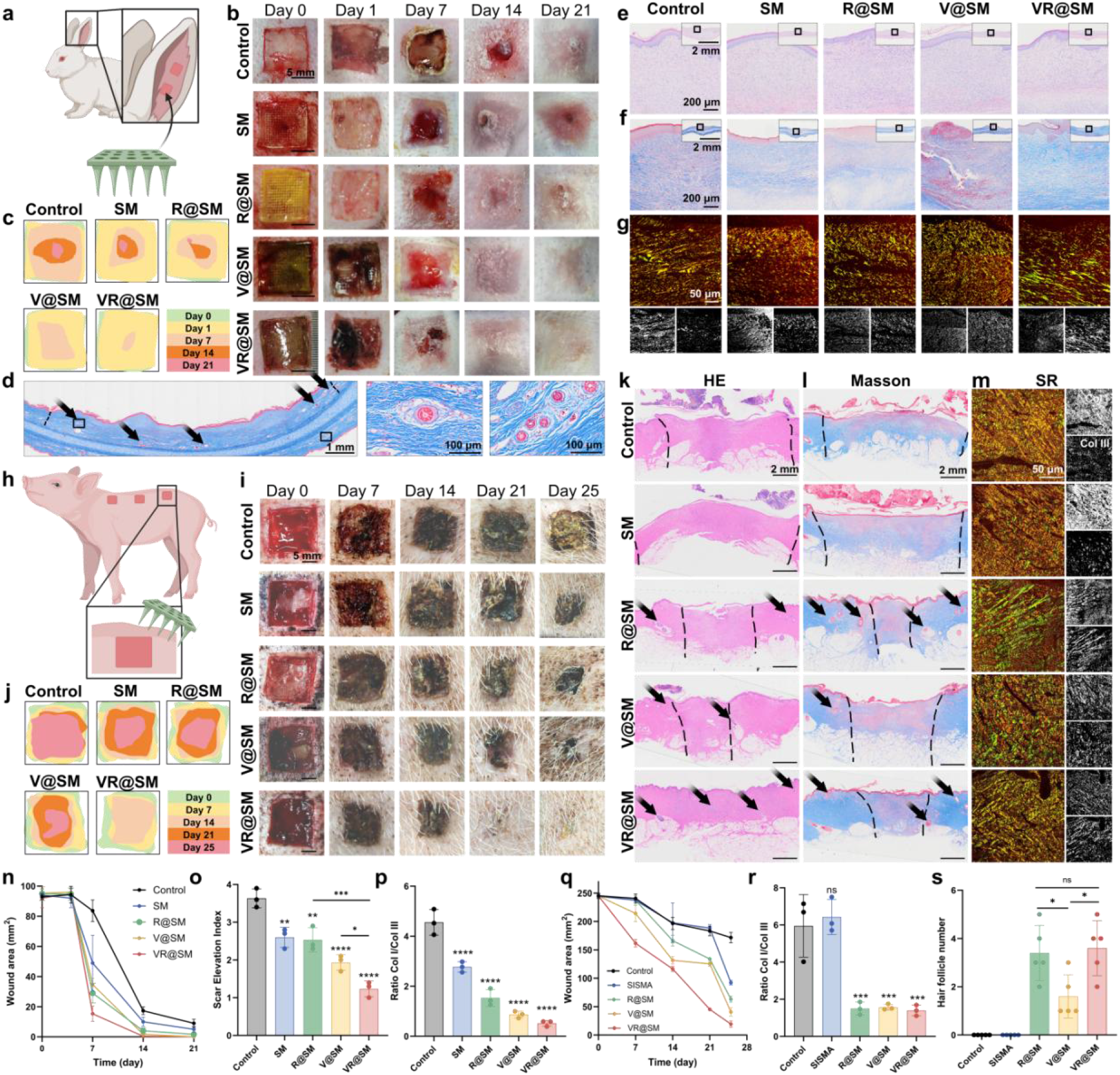
**VR@SM microneedles promote scar-suppressed skin regeneration in rabbit and porcine wounds. a–g, Rabbit-ear full-thickness wounds: experimental timeline (a), representative wound photographs (b), closure overlays (c), Masson’s trichrome staining showing follicular structures in regenerated tissue (d), and H&E, Masson’s trichrome and picrosirius-red staining of wound-centre sections on day 21 (e–g). h–m, Bama miniature-pig full-thickness wounds: experimental timeline (h), representative wound photographs (i), closure overlays (j), and H&E, Masson’s trichrome and picrosirius-red staining of wound-centre sections on day 25 (k– m). n–p, Rabbit wound closure (n; n=3), scar-elevation index (o; n=3), and type I/type III collagen ratio (p; n=3). q–s, Porcine wound closure (q; n=3), type I/type III collagen ratio (r; n=3), and regenerated follicle number in wound-centre sections (s; n=5). Data are mean ± s.d. One-way ANOVA with Dunnett’s multiple-comparison test was used for comparisons with untreated controls; selected pairwise comparisons used two-tailed unpaired Student’s t-tests. \**P*<0.05, \*\**P*<0.01, \*\*\**P*<0.001, \*\*\*\**P*<0.0001; ns, not significant**.

Hair shafts became visible within VR@SM-treated wound areas, and histological sections showed newly formed follicular structures within regenerated tissue (Fig. 2d). Haematoxylin and eosin, Masson’s trichrome and picrosirius-red staining showed less hypertrophic tissue accumulation and less collagen-dense remodelling after dual-cue treatment than in untreated wounds (Fig. 2e–g). Quantitatively, the scar-elevation index in the VR@SM group was reduced to 33.88% of the untreated control value. Collagen-area measurements and the type I/type III collagen ratio further supported reduced fibrotic remodelling, while regenerated follicle counts confirmed appendage formation in the wound centre (Fig. 2n–p).

Single-cue groups produced partial benefits, but neither consistently reproduced the combined closure, collagen-remodelling and follicle-regeneration phenotype of VR@SM. These results support the premise that early anti-fibrotic restraint and sustained pro-regenerative stimulation are complementary rather than interchangeable.

### The regenerative phenotype extends to miniature-pig skin

We next evaluated the same treatment logic in Bama miniature pigs. Multiple full-thickness wounds were generated on the dorsum and flank, and treatments were applied once immediately after wound creation. VR@SM-treated wounds showed accelerated closure and the most evident macroscopic restoration among the groups (Fig. 2h–j,q).

Histological analysis of the wound centre identified follicular structures in VR@SM-treated tissue and showed less collagen-dense remodelling by Masson’s trichrome and picrosirius-red staining (Fig. 2k–m). RA-containing treatments promoted hair regeneration, whereas the dual-cue VR@SM formulation provided the most balanced outcome across wound closure, collagen remodelling and follicle number (Fig. 2q–s). The detection of appendage-bearing repair in porcine skin argues that the regenerative phenotype is not restricted to rodent wound-induced hair neogenesis and can emerge in a large-animal skin environment.

## Discussion

This study supports a simple but consequential shift in regenerative design: controlled injury can be treated as a therapeutic input rather than merely a side effect of delivery. The microneedle array provides a spatially defined microtrauma pattern, while staged verteporfin and retinoic-acid exposure instructs the resulting repair niche. In both rabbit and miniature-pig wounds, this combination was associated with reduced scar-like collagen remodelling and the formation of follicular structures.

The findings build on prior evidence that wound geometry and mechanotransduction regulate the balance between scarring and regeneration [8–13,18–23]. Verteporfin-mediated restraint of YAP-associated signalling may limit early fibrotic reinforcement, whereas retinoic acid can support epithelial and dermal programmes associated with follicle induction [24–26]. The partial responses observed with single-cue formulations are consistent with a model in which removing a fibrotic barrier and providing a regenerative signal are both required for a robust outcome.

The large-animal result is particularly relevant because porcine skin more closely resembles human skin than murine skin in epidermal thickness, dermal architecture and wound-healing behaviour. Within these boundaries, the rabbit and porcine evidence establishes the priority of the core concept: a microneedle-induced repair window can be molecularly instructed to favour appendage-bearing skin restoration over fibrotic closure.

## Methods

### Synthesis of photocrosslinkable SISMA

Enzymatically digested and lyophilized porcine small intestinal submucosa (SIS) was reconstituted in ultrapure water at 0.5% (w/v). Methacrylic anhydride was added dropwise at a 1:20 molar ratio of native SIS free amino groups to methacrylic anhydride while the pH was maintained at 8.0. The reaction proceeded overnight at room temperature. The product was dialysed against ultrapure water using 14-kDa molecular-weight-cut-off tubing for 5 days and lyophilized. For photocrosslinking, SISMA was reconstituted at 10% (w/v) with 0.1% (w/v) lithium phenyl-2,4,6-trimethylbenzoylphosphinate and exposed to 365-nm ultraviolet light at 10 mW cm^-2^ for 30 s.

### Fabrication of layered microneedle patches

Native SIS solution (10% w/v) was mixed with verteporfin or retinoic acid to final concentrations of 0.21 mg ml− ^1^ and 3 mg ml^− 1^, respectively. For VR@SM patches, 100 μl of verteporfin-containing precursor was deposited into a polydimethylsiloxane negative mould containing a 15 × 15 array of 600-μm needles and centrifuged at 4,000 × g for 5 min before drying. SISMA precursor containing 0.1% photoinitiator was then added, centrifuged, dried and photocrosslinked. Finally, 10 μl of retinoic-acid-containing SIS solution was deposited onto the backing and dried. Unloaded SM, V@SM and R@SM patches were fabricated using the corresponding omission or single-cue formulation. Patches were stored protected from light at 4 °C under dry conditions.

### Animals, anaesthesia and treatment groups

Male New Zealand white rabbits (6–8 weeks old; 2.0–2.5 kg) and Bama miniature pigs (2–3 months old; approximately 20 kg) were housed under standard conditions with a 12-h light/dark cycle, 22 °C ambient temperature and 50–60% humidity. All operations were conducted under general anaesthesia with perioperative analgesia. Rabbits received intramuscular ketamine (35 mg kg^-1^) and xylazine (5 mg kg^-1^). Pigs were premedicated intramuscularly, intubated and maintained with 1.5–2.5% isoflurane. Surgical sites received local infiltration with 0.25% bupivacaine.

Experimental groups were untreated control, unloaded SM, V@SM, R@SM and VR@SM. Microneedle patches were applied once immediately after wound creation and were not replaced during follow-up.

### Rabbit-ear and porcine full-thickness wound models

Four 10 × 10-mm full-thickness wounds were created on the ventral surface of each rabbit ear by excision to the perichondrial surface while preserving cartilage. In Bama miniature pigs, multiple 1.5 × 1.5-cm full-thickness wounds were created on the dorsum and flank to the layer immediately above subcutaneous fat. Pig wounds were covered with sterile petrolatum gauze and elastic bandages, and animals wore protective jackets. Wounds were photographed at prespecified time points. Rabbit tissues were collected on day 21 and porcine tissues on day 25 after euthanasia by anaesthetic overdose.

### Histology and image analysis

Tissues were fixed in 4% paraformaldehyde, paraffin embedded and sectioned at 5 μm. H&E staining was used to assess epithelialization, tissue architecture and appendage formation. Masson’s trichrome staining was used to assess collagen deposition, and picrosirius-red staining was examined under polarized light to distinguish thick type I collagen from thin type III collagen. Whole-slide images were acquired using an Olympus VS200 scanner. Wound area, scar-elevation index, collagen area, type I/type III collagen ratio and follicle number in central wound sections were quantified using ImageJ and OlyVIA.

### Statistical analysis

Analyses were performed using GraphPad Prism 10. Data are presented as mean ± s.d. For comparisons involving more than two groups, one-way ANOVA followed by Dunnett’s multiple-comparison test was used. Selected pairwise comparisons were assessed using two-tailed unpaired Student’s t-tests. *P*<0.05 was considered statistically significant.

### Ethics statement

All animal experiments were conducted in accordance with institutional guidelines and relevant regulations after approval by the responsible institutional animal care and use committee.

## Data availability

The data summarized in the two figures are available from the corresponding authors upon reasonable request. Expanded source data and additional material, mechanistic and transcriptomic datasets are being prepared for release with the complete study. No public accession number is claimed in this concise version.

## Author contributions

K.W. and Z.-Y.F. contributed equally. K.W. designed the study, developed animal models, performed in vivo efficacy evaluation and conducted mechanistic analyses. Z.-Y.F. performed cell-based studies and designed and characterized the biomaterials and microneedle patches. Z.-Y.Z., Q.-F.L. and H.-Q.X. provided supervision, funding support and input into study design. All authors reviewed and approved the manuscript.

## Competing interests

The authors declare that intellectual-property applications related to SISMA and/or components of the described microneedle platform may exist. The authors declare no other competing interests.

## Notes

### Competing Interest Statement

The authors have declared no competing interest.

